# HIC1 interacts with FOXP3 multi protein complex: a novel mechanism to regulate human regulatory T cell differentiation and function

**DOI:** 10.1101/2023.05.15.540505

**Authors:** Syed Bilal Ahmad Andrabi, Kedar Batkulwar, Santosh D. Bhosale, Robert Moulder, Meraj Hasan Khan, Tanja Buchacher, Mohd Moin Khan, Ilona Arnkil, Omid Rasool, Alexander Marson, Ubaid Ullah Kalim, Riitta Lahesmaa

## Abstract

Transcriptional repressor, hypermethylated in cancer 1 (HIC1) participates in a range of important biological processes, such as tumor repression, immune suppression, embryonic development and epigenetic gene regulation. Further to these, we previously demonstrated that HIC1 provides a significant contribution to the function and development of regulatory T (Treg) cells. However, the mechanism by which it regulates these processes was not apparent. To address this question, we used affinity-purification mass spectrometry to characterize the HIC1 Interactome in human Treg cells. Altogether 61 high-confidence interactors were identified, including IKZF3, which is a key transcription factor in the development of Treg cells. The biological processes associated with these interacting proteins include protein transport, mRNA processing, non-coding (ncRNA) transcription and RNA metabolism. The results revealed that HIC1 is part of a FOXP3-RUNX1-CBFB protein complex that regulates Treg signature genes thus improving our understanding of HIC1 function during early Treg cell differentiation.

**Highlights:**

- Systematic characterization of HIC1 interactome in regulatory T cells by Affinity Purification-Mass Spectrometry
- HIC1 binds to the *RUNX1* promoter and regulates its expression
- HIC1-a part of FOXP3-RUNX1-CBFB transcriptional complex

**Graphical abstract:** 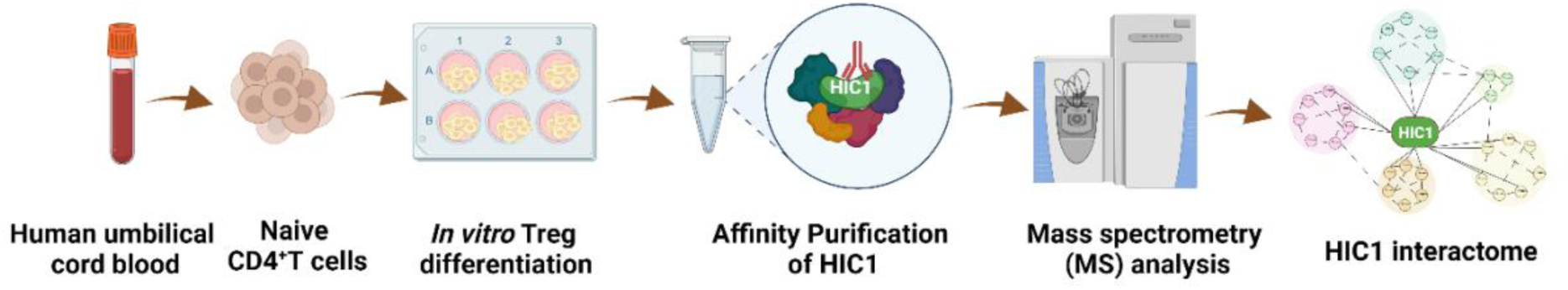

## 1. Introduction

Regulation of the immune system is critical for controlling autoimmunity and fighting cancer. [1]. Central to these processes, regulatory T (Treg) cells maintain immune tolerance and the balance between pro-and anti-inflammatory responses [2]. The majority of Treg cells are produced in the thymus (tTreg), while some acquire regulatory phenotype in the periphery (pTreg) [3]. Treg cells can also be induced *in vitro* (iTreg) from naïve CD4^+^ T cells through their activation in the presence of Treg cell polarizing cytokines [4]. The lineage-specific transcription factor (TF) FOXP3 is essential for Treg differentiation, stability, and function [4,5]. Besides FOXP3, several other TFs, e.g., IKZFs, NR4As, C-REL, NFAT, SMAD factors, STAT5, and RUNX1 act in concert and play an important role in Treg cell differentiation [6–13].

Recently, we characterized the role of a novel TF hypermethylated in cancer 1 (HIC1), in human iTreg cell differentiation and showed that it contributes to suppressive function and lineage specification of Treg cells without influencing expression of FOXP3 [14]. It is a member of the Kruppel/Zinc Finger and BTB (POK/ZBTB) protein family and is downregulated in a variety cancers [15]. In iTreg cells, HIC1 deficiency led to a considerable loss of suppressive capability with a concomitant increase in the expression of effector T cell associated genes including GATA3, TBX21 and IFNG [14]. However, the molecular mechanisms by which HIC1 regulates the suppressive capacity of iTreg cells, were lacking. As a follow up, to address this deficit, we comprehensively studied the protein interaction network of HIC1 and validated the key interactors. The results indicate that HIC1 is a vital part of a protein complex that regulates Treg signature genes. This study sheds light on the mechanism of HIC1 action during human Treg cell differentiation.

## 2. Materials and methods

### 2.1 CD4^+^ cell isolation and differentiation to iTreg Cells

CD4^+^ T cells were isolated from human umbilical cord blood as described previously [14]. Briefly, umbilical cord blood was layered on Ficoll (GE Healthcare, Cat# 17-1440-03) to isolate white blood cells and the Dynal bead CD4^+^ isolation kit (Invitrogen, Cat# 11331D) was used to isolate CD4^+^ T cells. CD25 MicroBeads II and LD columns (Miltenyi Biotec, Cat# 130-092-985 and Cat# 130042901, respectively) were used for selection of CD25^−^ T-cells. CD4^+^ CD25^−^ cells from multiple donors were pooled before activation. For iTreg cultures, cells were activated with plate-bound anti-CD3 (500 ng/one well of 24-well culture plate; Beckman Coulter, REF# IM-1304) and soluble anti-CD28 (500 ng/ml; Beckman Coulter, REF# IM1376) antibodies in presence of cytokines i.e., IL-2 (12 ng/ml; R&D Systems); all-trans retinoic acid (ATRA) (10 nM; Sigma-Aldrich); TGF-β1 (10 ng/ml; R&D Systems) and human serum (10%) and cultured for 72 h at 37 °C in 5% CO_2_.

### 2.2 Co-immunoprecipitation (Co-IP)

HIC1 immunoprecipitation (IP) was performed using anti-HIC1 antibody (Santa Cruz Biotechnology, Cat# sc-271499) that recognizes 611-733 C-terminus region of human HIC1, together with the respective control IgG antibody, in three biological replicates. Similarly, reverse IPs were performed with anti-RUNX1 antibody (Santa Cruz Biotechnology, Cat# sc-365644), anti-FOXP3 antibody (eBioscience, Cat# 14-4776-82), anti-IKZF3 antibody (Abcam, Cat# ab139408) and anti-CBFB antibody (Abcam, Cat # 62184). The Pierce™ MS-Compatible Magnetic IP Kit (ThermoFisher Scientific, Cat# 90409) was used. The respective Immunoglobulin G (IgGs) were used in control IPs. Briefly, differentiated iTreg cells were harvested on ice and washed with PBS (2X). Cells were then re-suspended in ice-cold IP-MS Cell Lysis Buffer, followed by incubation on ice for 20 min with periodic mixing as recommended by the manufacturer. The protein lysate was centrifuged at ∼13,000 g for 10 min to pellet down the cell debris. The supernatant was transferred to a new microcentrifuge tube and protein concentration was determined by using a DC protein assay kit (BioRad, Cat# 500-0116). Cell lysates (500-1000 μg) were mixed with antibodies (1:50 dilution) and the antibody/lysate solution was diluted with the IP-MS cell lysis buffer and incubated overnight at +4 °C to form the immune complex, according to the manufacturers protocol. On the next day, the immune complex was incubated with Protein A/G magnetic beads for 1 h at room temperature (RT) followed by washing the beads and elution. The eluate containing the target antigen was then transferred to a new low protein-binding microcentrifuge tube and dried by vacuum concentrator prior to MS sample preparation and western blot.

### 2.3 Western blotting

Cells were resuspended in RIPA buffer (ThermoFisher, Cat# 89901) and sonicated on ice for 5-10 min using a Bioruptor® sonicator (Diagenode). The cell lysate was centrifuged at high speed (16,000 g) and protein containing supernatant was transferred to a new tube. Protein concentration was determined using a DC Protein assay (Bio-Rad, Cat# 500-0116). Equal amounts of protein were loaded onto acrylamide gel (Bio-Rad Mini PROTEAN® TGX precast gels). For the transfer of proteins to the PVDF membrane, mini transfer packs from Bio-Rad were used. Primary/secondary antibody incubations were performed in 5% Bovine serum albumin (BSA) in TBST buffer (0.1% Tween 20). The following antibodies were used, RUNX1 (Santa Cruz Biotechnology, Cat# sc-365644); CBFB antibody (Abcam, Cat # 62184); HIC1 (Santa Cruz Biotechnology, Cat# sc-271499); IKZF3 (Abcam, Cat# ab139408) FOXP3 (eBioscience, Cat# 14-4776-82) and Beta-actin (Sigma, cat # A5441).

### 2.4 Sample preparation for mass-spectrometry analysis

The immunoprecipitated proteins (from both the HIC1 and IgG control bait) were digested with trypsin. Briefly, urea buffer (8 M urea, 50mM Tris-HCl pH 8.0) was added to denature the proteins, followed by addition of dithiothreitol to a final concentration of 10 mM and incubation at 37 °C for 1 h for reduction. The reduced disulphide bridges were subsequently alkylated using iodoacetamide (∼14 mM) at RT for 30 min in dark condition. The samples were diluted to a urea concentration of less than 1 M and digested with sequencing grade modified trypsin (0.29 µg per sample) at 37 °C overnight (16 h). The tryptic digests were acidified and desalted using in house made C18 Stage Tips (3M, Cat No 2215). The eluates from the Stage Tips were dried in a vacuum centrifuge (Thermo Fisher Scientific) and stored at −80 °C until further analysis.

### 2.5 LC-MS/MS analysis

The desalted tryptic peptides were reconstituted in formic acid/acetonitrile mixture and the peptide amounts were estimated using a NanoDrop-1000 UV spectrophotometer (Thermo Fisher Scientific). The samples were analysed by LC-MS/MS using an Easy-nLC 1200 coupled to Q Exactive HF mass spectrometer (Thermo Fisher Scientific). A 20 x 0.1 mm i.d. Pre-column coupled with a 75 µm x 150 mm analytical column (both in-house packed with 5 µm Reprosil C18; Dr Maisch GmbH) were used for sample loading and separation, respectively. Peptides were eluted using a gradient from 5 to 36% B in 50 min at a flow rate of 300 nl/min. Tandem mass spectra were acquired in positive ion mode using HCD fragmentation of the 15 most intense ions from each precursor scan. The Orbitrap resolution was set to 120,000 at m/z 200 for the full scan MS spectra, with a maximum injection time of 100 ms and a target value of 3 x 10^6^ in the 300-1650 m/z range. The tandem mass spectra were acquired at a resolution of 15,000 at 200 m/z with a target value of 5 x 10^4^ and a maximum injection time of 150 ms. The lowest fixed first mass of 120 m/z was used and to the repeatedly identified peptides were excluded for 20 s. The samples were analyzed in triplicate in randomized batches.

### 2.6 Data analysis

The mass spectrometry (MS) raw files were analyzed using MaxQuant software version 1.6.0.16 [19] and searched against a human UniProt FASTA sequence database (downloaded, May 2019 and containing 20415 entries) with common contaminants using the Andromeda search engine [20]. The specified search criteria were for trypsin digestion with up to two missed cleavages, a fixed carbamidomethyl modification of cysteine residues and variable modifications of methionine oxidation and N-terminal acetylation. The false discovery rate (FDR) was set to 1% at the peptide and protein level.

The ‘proteinGroup.txt’ table generated from the search was filtered to remove contaminants, proteins only identified by site and reverse hits using Perseus 1.6.2.3 [16]. Proteins identified with two or more unique peptides were retained. The protein LFQ values were log2 and the median values calculated for the technical replicates. The data was further filtered to remove the influence of inconsistently detected features, based on the inclusion criteria of three valid values in at least one group, i.e. IgG or HIC1. For statistical analysis of the data, the mass spectrometry interaction statistics (MiST) algorithm [17]was used. The algorithm compares the results from IgG control to pulldowns bait and then computes a score for each protein. The MiST scoring algorithm uses the protein signal intensity,reproducibility and itsspecificity to the bait, to calculate a score ranging from 0 to 1. The criteria of a MiST score, ≥ 0.75 with the HIC1 bait and ≤ 0.75 with IgG control bait was applied to define the putative protein interactors [18]. Further filtering was made relative to an in-house database of proteins frequently detected with IgG baits in similar experiments (i.e. human Th cells with Dynabeads). Proteins detected in more than 70% of the previous measurements were excluded from the analysis. Cytoscape was used to build the protein-protein interaction (PPI) network of the identified interactors, combining existing PPI data from the STRING database [19,20]. The mass spectrometry proteomics raw data have been deposited to the ProteomeXchange Consortium via the PRIDE [16] partner repository with the dataset identifier PXD039337.

### 2.7 Selected reaction monitoring mass spectrometry (SRM-MS)

Selected reaction monitoring mass spectrometry was used to validate the relative abundance of BCOR, FOXP3, RUNX1, CBFB, IKZF3 and HIC1 in immuno-precipitates from iTreg cells. ACTB and GAPDH were also measured as a reference protein. Heavy-labeled synthetic peptides (lysine ^13^C_6_ ^15^N_2_ and arginine ^13^C_6_ ^15^N_4_) were obtained for the targets of interest (PEPotec, Grade 2, Thermo Fischer Scientific). For these validations, IP’s were prepared from the differentiated iTreg cells from three donors. Skyline software[21] was used to develop the SRM method, check the peak integration, normalize the data and for statistical analysis.

The samples were prepared using the same digestion and desalting protocols used for discovery. These were then spiked with synthetic heavy labelled analogues of the peptide targets and a retention time standard (MSRT1, Sigma) for scheduled SRM. The LC-MS/MS analyses were conducted using Easy-nLC 1000 liquid chromatography (Thermo Scientific) coupled to a TSQ Vantage Triple Quadrupole mass spectrometer (Thermo Scientific). The column configuration included a 20 x 0.1 mm i.d. pre-column in conjunction with a 150 mm x 75 µm i.d. analytical column, both packed with 5 µm Reprosil C18-bonded silica (Dr Maisch GmbH). A separation gradient was used from 5% to 21% B in 11 min, then to 36% B in 9 min, to 100% in 2 min, then ending with an 8 min isocratic period. A flow rate of 300 nl/min was used, with the mobile phase compositions as indicate above. The raw SRM data are available through Panorama (https://doi.org/10.1074/mcp.RA117.000543) with the dataset identifier PXD038532. The estimated injected amount was 250 ng of endogenous sample, spiked with 50 fmol of heavy labelled peptides. Measured peptides used are listed in Table S1.

### 2.8 Proximity ligation assay (PLA)

PLA assay was performed following the manufacturer’s protocol (Duolink®PLA, Sigma). The Treg cells were fixed with 4% paraformaldehyde for 15 min at RT and then plated on Poly-L-lysine coated (10 μg/ml) coverslips using a cytospin for 5 min at 800 RPM. The cells were permeabilized for 10 min with PBS containing 0.5% Triton X-100 at RT. A blocking solution was added to the cells for 30 min at 37 °C, followed by incubation with primary antibodies (in blocking solution) anti-HIC1 (Santa Cruz Biotechnology, Cat# sc-271499), anti-IKZF3 (Abcam, Cat# ab139408), anti-CBFB (Cell Signaling Technology, Cat# 62184), anti-FOXP3 (Invitrogen, Cat# PAI-806), RUNX1 (Santa Cruz Biotechnology, Cat# sc-365644) anti-GFP (mouse) (Abcam, Cat# ab1218) and anti-GFP (rabbit) (Invitrogen Cat# A11122). After washing with buffer A, PLA probes were incubated for 1 h at 37 °C, followed by a ligase reaction step performed for 30 min at 37 °C. In the final step, polymerase solution was added and the cells were incubated for 100 min at 37 °C for the amplification. All the incubations were performed in a preheated humidity chamber. Post-amplification, the coverslips were washed with buffer B from the PLA kit and mounted with Vectashield having DAPI (Vector Laboratories). The PLA signal was detected by using a confocal microscope 3iCSU-W1 spinning disc microscope equipped with a 100x (NA 1.4 oil, Plan-Apochromat, M27) objective and Evolve 512 EMCCD camera (Photometrics). PLA signals per cell were calculated by dividing the amount of PLA signal dots in one field of view, determined using Cell Profiler software [22].

### 2.9 CRISPR-Cas9 mediated HIC1 ablation

The in vitro assembly of guide RNA (gRNA) with the Cas9 protein was carried out as described previously [23]. Briefly, 80 μM of gRNA reagents were prepared by combining equimolar amounts (1:1) of HIC1 targeting (5’-GCATGACAACCTGCTCAACC-3’) or non-targeting control (NT) (5’-CGTTAATCGCGTATAATACG-3’) crisprRNA (crRNA) with trans-activating crispr RNA (tracrRNA) scaffold (both synthesized by IDT) followed by incubation at 37 °C for 30 min. Assembled 80 μM gRNA was then mixed with equal volume of 40 μM recombinant *S. pyogenes* Cas9-nuclear localization sequence (NLS) purified protein (QB3 Macro Lab, University of California, Berkeley) (2:1 gRNA to Cas9 molar ratio) and 1μl of 100 μM Ultramer DNA oligonucleotide enhancer (IDT), followed by incubation for 10 min at 37 °C for a final concentration of 20 μM CRISPR-Cas9 ribonucleoprotein (RNP). Freshly purified CD4^+^ CD25^−^ cells, resuspended in Opti-MEM™ (Gibco by Life Technologies, Cat# 31985-047), were transfected with RNP complexes using Nucleofector 2C system (Lonza) (4 × 10^6^ cells per cuvette in 100 μL of Opti-MEM using U-014 nucleofection program). One milliliter of pre-warmed culture media was added immediately after nucleofection, and cells were then transferred into a 6-well plate, and additional culture media was added to a final volume of 3ml. After nucleofection, cells were rested for 24 h in RPMI supplemented with 10% serum, followed by cell culturing under iTreg condition for 72 h, as described above.

### 2.10 ChIPmentation-qPCR analysis

Tagmentation-based chromatin immunoprecipitation (ChIPmentation) was conducted as described previously [24]. Briefly, fixed chromatin from differentiated Treg cells (three million cells) was sonicated and immunoprecipitated with anti-HIC1 or control IgG antibody. Genomic DNA was probed by qPCR for *RUNX1* promoter region. PCR primers for ChIPmentation are listed in Table S2. PCR primers were designed by LightCycler® Probe Design Software from Roche. Real-time PCR was performed using custom TaqMan Gene Expression Assay reagent on QuantStudio 12K Flex Real-Time PCR System (Thermo Scientific).

### 2.11 Luciferase Assay

HIC1 binding on *RUNX1* promoter was assessed using Dual-Luciferase® Reporter Assay by cloning the ChIP-defined genomic region upstream of a minimal promoter driving a luciferase gene (pGL4.10 [luc2/minP]; Promega). Wild type HIC1 binding motif of the *RUNX1* proximal promoter (5′-TGCCCTGG-3′) or the corresponding mutant target sequence (ΔRunx1: 5′-AAAAATGG-3) was cloned upstream of a minimal promoter in a luciferase reporter plasmid pGL4.10.

Näive CD4^+^ T cells were nucleofected with Runx1-pGL4minP or ΔRunx1-pGL4minP or empty pGL4minP construct and renilla luciferase plasmid [25]. Post nucleofection, cells were rested and cultured under Treg polarizing condition for 72 h, after which cells were collected and luciferase assay was performed using the Dual Luciferase Reagents (Promega). Firefly luciferase activity was normalized to renilla luciferase activity for each sample and expressed as fold change over empty pGL4-minP. Δ*RUNX1*-pGL4minP with mutated promoter region served as a negative control.

### 2.12 Ethical approval

Usage of the blood of unknown donors is approved by the Ethics Committee, Hospital District of Southwest Finland.

## 3. Results

### 3.1 Identification of HIC1 protein interaction network in iTreg cells using affinity purification–mass spectrometry

To better understand how HIC1 regulates the development and function of human iTreg cells, we performed HIC1 immunoprecipitation (IP) (Figure S1) followed by mass spectrometry (MS) to identify the interacting partners of HIC1. We identified 61 proteins as high confidence HIC1 protein interactors (Figure 1A; Table S3). Notably, RUNX1, CBFB, and IKZF3, which have important roles in Treg cell development and function, were detected as components of this interactome [26–29]. Additionally, many tRNA synthetases (EPRS, DARS, IARS, QARS, RARS, MARS, KARS and LARS) and tRNA synthetase complex interacting multifunctional proteins (AMP1 and AMP2) were identified as HIC1 interactors (Figure 1A; Table S3). Interestingly, many of the interacting proteins are associated with RNA regulatory processes. For instance RPRD1B and RPRD2 are known to preferentially bind to the phosphorylated CTD of RNAP II, leading to decreased Ser-5 and Ser-7 phosphorylation of RNAP II at target gene promoters and regulate gene transcription [30]. Likewise, PARP13 and splicing factor SRSF1 were also identified in the HIC1 interactome and are reported to function in the regulation of RNA stability and splicing [31,32]. Similarly, we identified association of HIC1 with E3

**Figure 1:**
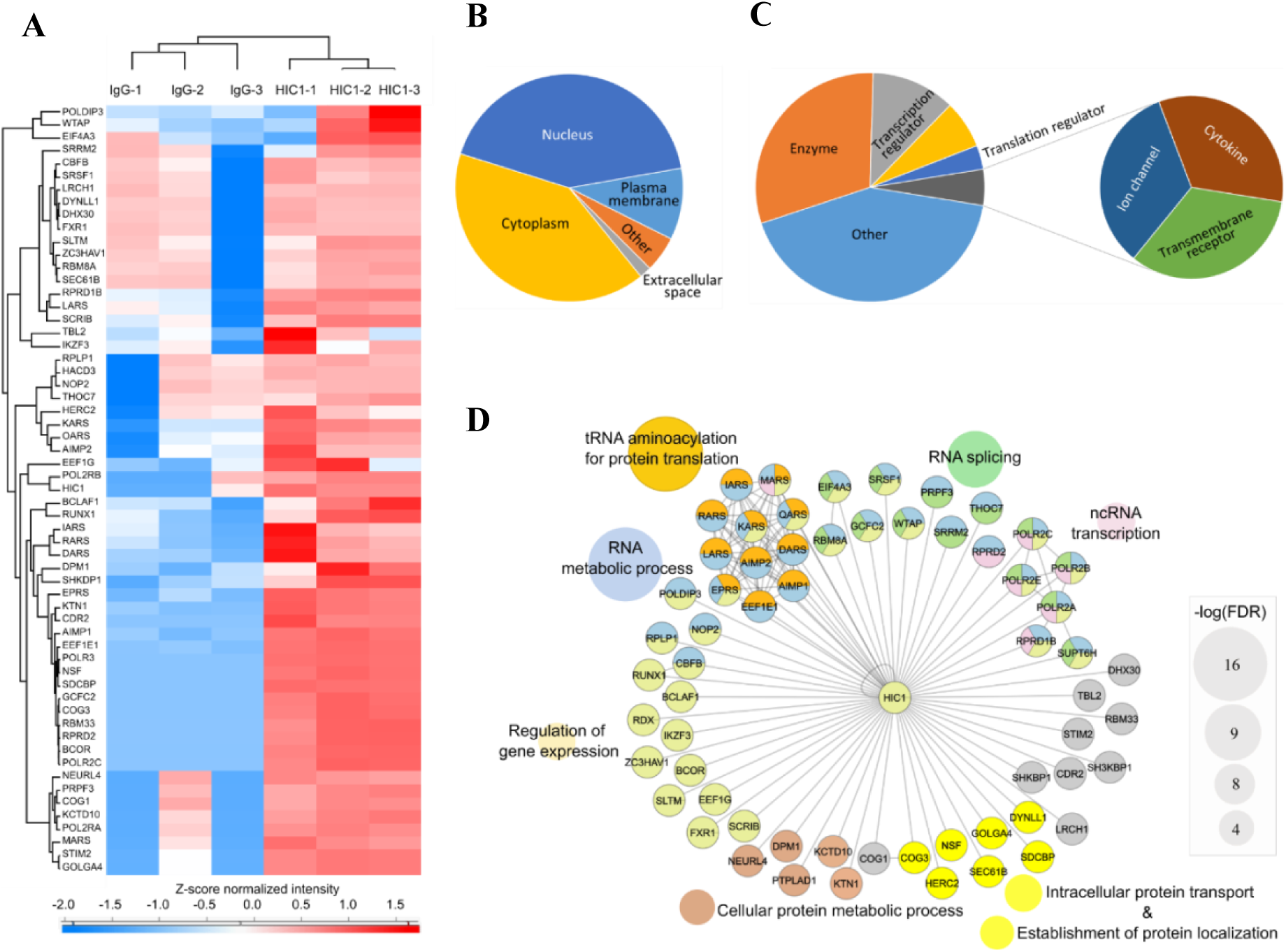
The HIC1 Interactome. Following affinity purification and mass spectrometry (AP-MS), statistical analysis of interactions using the MIST algorithm and filtering for common contaminants, a panel of interacting proteins was identified. The proteins are shown as a heat-map (**A)**, clustered on the basis of their Z-score normalized LFQ intensities from the bait and control measurements. Representations of their cellular localization (**B**) and functional classes (**C**) are shown as pie charts. The interactors are further represented as an interaction network (**D**), with the enriched biological processes indicated by the use of colored circles and strength of the enrichment indicated by the size of the peripheral circles. Further representation of this data is provided in **Table S4.**

Ubiquitin-Protein Ligase HERC2, a member of the HECT family of E3 ubiquitin-protein ligases implicated in DNA damage repair responses [33]. Furthermore, BCOR (BCL-6 interacting corepressor) was among the detected HIC1 interacting proteins (Figure 1A; Table S3). BCOR is known to interact with the BTB/POZ domain that is a specific feature of a small subset of zinc finger proteins including BCL-6 and HIC1 [34].

Ingenuity Pathways Analysis (IPA; Qiagen) was used to summarize the cellular locations of the HIC1 interactors. The analysis revealed that the majority of the HIC1 interactors were associated with nucleus, followed by the cytoplasm, whilst small fraction of interactors were distributed in plasma membrane and extracellular space (Figure 1B). Although HIC1 is primarily localized in nucleus[14], it might interact with cytoplasmic proteins after they are translocated to nucleus or when HIC1 is in the cytoplasm at different stages of cell activation and differentiation. Enzymes was the major functional class of the HIC1 interacting proteins followed by transcriptional and translational regulators (Figure 1C). Additionally, the other functions associated with HIC1 interacting proteins were related to cytokines, ion channels and transmembrane receptor proteins (Figure 1C). Gene ontology (GO) enrichment analysis on HIC1 interacting proteins revealed that most enriched biological processes were associated with regulation of gene expression, tRNA amino-acetylation for protein translation, RNA splicing, ncRNA transcription, RNA metabolism, cellular and metabolic process and protein transport (Figure 1D).

### 3.2 Validation of the HIC1 protein interactors in iTreg cells

For validation of the HIC1 interactome results, three different independent approaches, were used namely: selected reaction monitoring-mass spectrometry (SRM-MS), proximity ligation assay (PLA) combined with confocal microscopy, and co-immunoprecipitation (Co-IP)/reverse Co-IP followed by western blotting. For this, we selected CBFB, RUNX1, and IKZF3 since they are known to play important role in Treg cell function and homeostasis [26,27,29,35]. Further, we also included BCOR for SRM validation as it was among the top HIC1 interactor and is known to play role in maintaining the lineage stability and suppressive function of Treg cells [36]. Additionally, although FOXP3 was not among the identified HIC1 interacting proteins, we included it in the validation analysis, as it is a well-known interactor of both RUNX1-CBFB protein complex and IKZF3 [29,37]. As illustrated in Figure 2A, the SRM-MS analysis confirmed HIC1 interaction with CBFB, RUNX1, IKZF3, BCOR and FOXP3. The interaction of HIC1 with RUNX1, IKZF3, CBFB and FOXP3 was further confirmed with Co-IP and PLA-confocal microscopy analysis (Figure 2 B-D).

**Figure 2:**
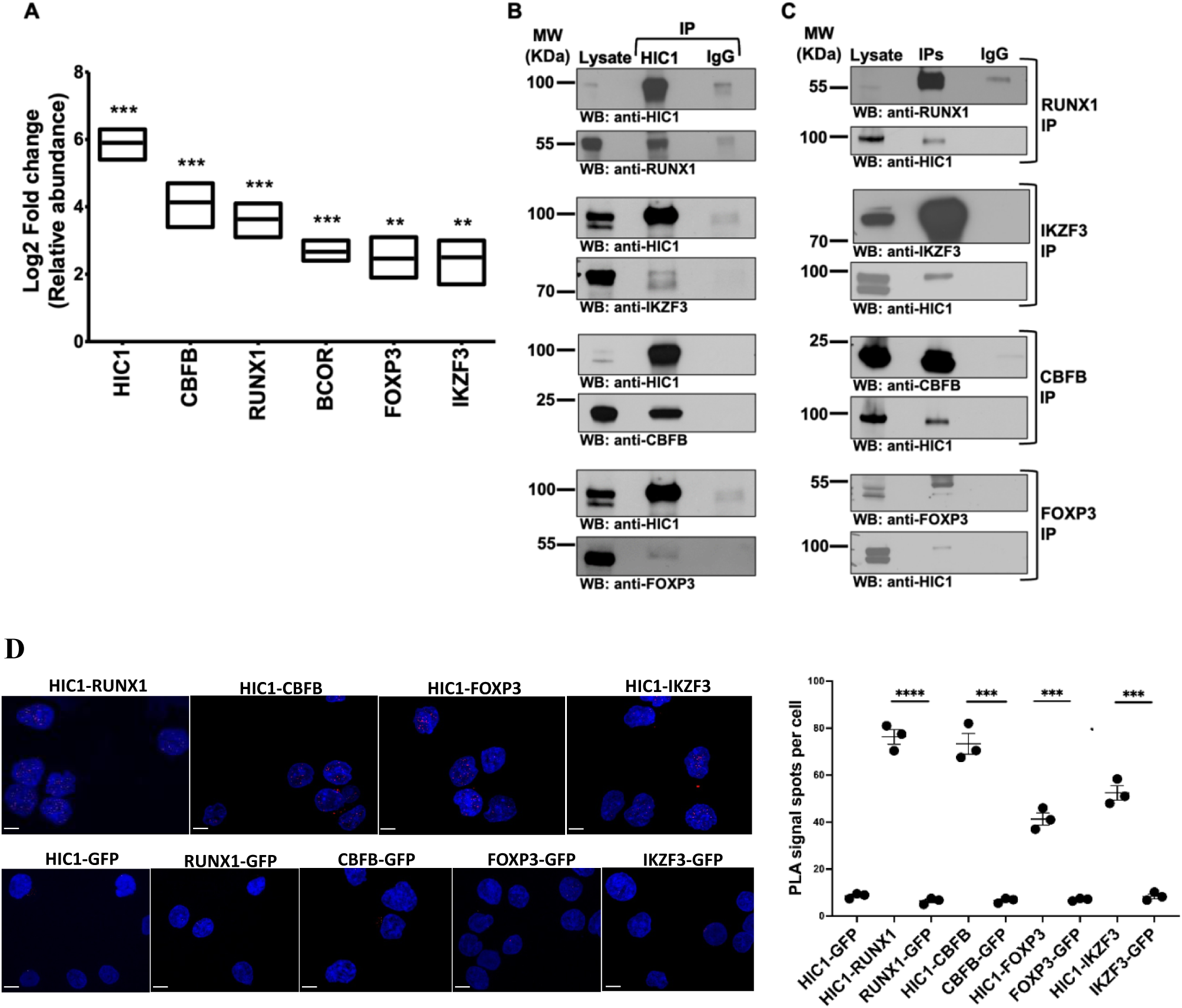
HIC1 interactome validation by SRM-MS, IP-WB, reverse IP-WB and PLA assay. (**A**) SRM-MS validation of the candidate HIC1 interacting partners. HIC1 IP was performed using HIC1 antibody in 72 h polarized iTreg cells and six proteins were validated by targeted proteomics. Averaged results from three replicates are presented in the form of a box plot. Statistical analysis was made by a two-tailed paired student’s T-test, **p-value represents < 0.005; and ***p-value represents < 0.0005. (**B**) Validation of HIC1 protein interaction by IP-western blot. HIC1 IP followed by WB detection with RUNX1, IKZF3, CBFB and FOXP3 antibodies was performed to validate the HIC1 interaction with RUNX1, IKZF3, CBFB and FOXP3. Total cell lysate (Input), control IP (IgG) and HIC1 IP are shown in the blots. A conformational specific secondary antibody was used to probe proteins without interference from the denatured IgG heavy (50 kDa) and light chains (25 kDa). A representative WB of three biological replicates is shown. (**C**) RUNX1-IP, IKZF3-IP, CBFB-IP and FOXP3-IP were performed using iTreg cell lysate followed by WB detection with HIC1 antibody to validate the HIC1 interaction with RUNX1, IKZF3, CBFB and FOXP3. A representative western blot of three biological replicates is shown. (**D**) Proximity ligation assay (PLA) with indicated antibody pairs in 72 h polarized iTreg cells. The GFP antibody was used as a negative control. DAPI was used to stain the nuclei. The scale bar is 7 μm. The graph shows the average number of PLA signals (spots) in each experiment (n=10 images from a representative experiment of n=3 experiments). Statistics by paired two-tailed T-test, where * ** represents p<0.01; and **** represents p<0.001.

### 3.3 HIC1 binds to RUNX1 promoter and modulates its expression

In our earlier HIC1 Chromatin Immunoprecipitation-sequencing (ChIP-seq) analysis [14], HIC1 binding was observed at the *RUNX1* promoter region in iTreg cells (Figure 3A). To verify the binding of HIC1 to *RUNX1* promoter region, we performed ChIPmentation assay followed by PCR. Higher fold enrichment of HIC1 at the *RUNX1* promoter compared to control IgG antibody was observed (Figure 3B), thus confirming the binding of HIC1 to the *RUNX1* promoter.

**Figure 3:**
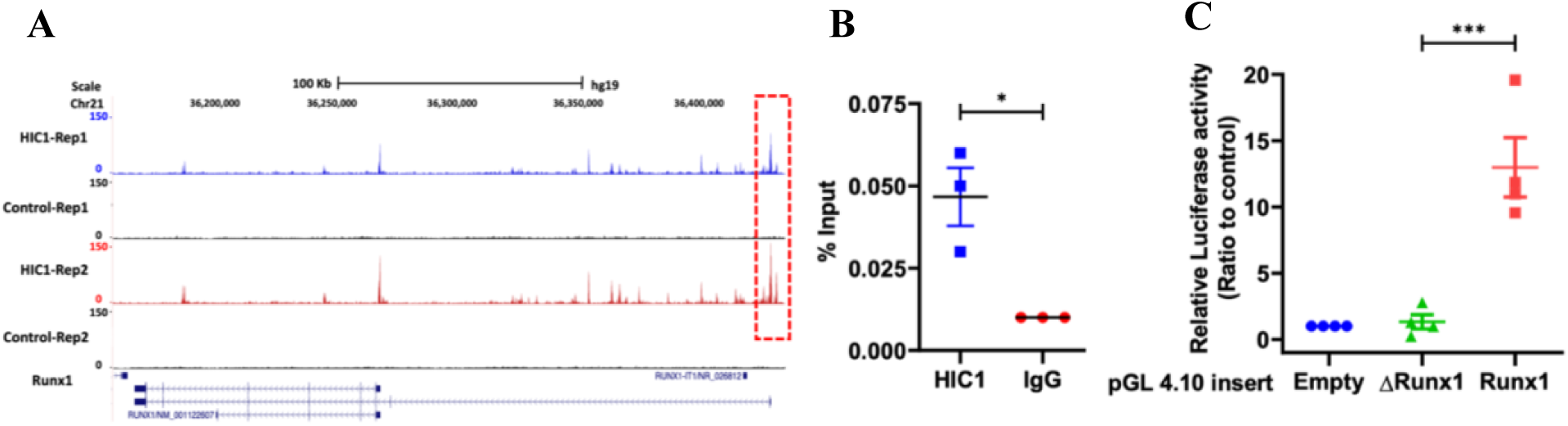
ChIPseq, Luciferase and ChIP-qPCR assays demonstrate the recruitment of HIC1 to the *RUNX1* promoter in iTreg cells. (**A**) ChIP-Seq analysis identifies binding sites of HIC1 at *RUNX1* promoter region in iTreg cells. UCSC genome browser snap shots for the *RUNX1* gene that demonstrates the HIC1 binding peaks as indicated in enclosed box. (**B**) ChIP-qPCR assay was performed in iTreg cells using anti-HIC1 IgG (HIC1) and mouse IgG (IgG) as control. Input of sheared chromatin was prepared prior to immunoprecipitation. (**C**) Relative luciferase activities were measured after co-transfection of *RUNX1* promoter (Runx1) or mutated Runx1 promoter (ΔRunx1) constructs along with the pRL-TK expression plasmid into iTreg cells. Statistical significance calculated using Student’s t-test (two-tailed paired) where ***denotes p value <0.0005. *denotes p value <0.05.

The observed binding of HIC1 to the *RUNX1* promoter suggested that it might be directly involved in the control of RUNX1 transcription in iTreg cells. To validate this hypothesis, a luciferase reporter assay, was used. For these analyses, the wild-type (WT) or mutated binding region of HIC1 at the *RUNX1* promoter was cloned upstream of firefly luciferase.

As shown in Figure 3C, the luciferase activity was significantly higher with WT *RUNX1* compared to mutated (Δ*RUNX1*) promoter construct. These findings suggest that HIC1 regulates transcriptional activation of *RUNX1* by binding to its promoter region. To study this further, CRISPR-Cas9-mediated HIC1 ablation in iTreg cells was performed which led to reduced RUNX1 expression. (Figure 4A-C).

**Figure 4:**
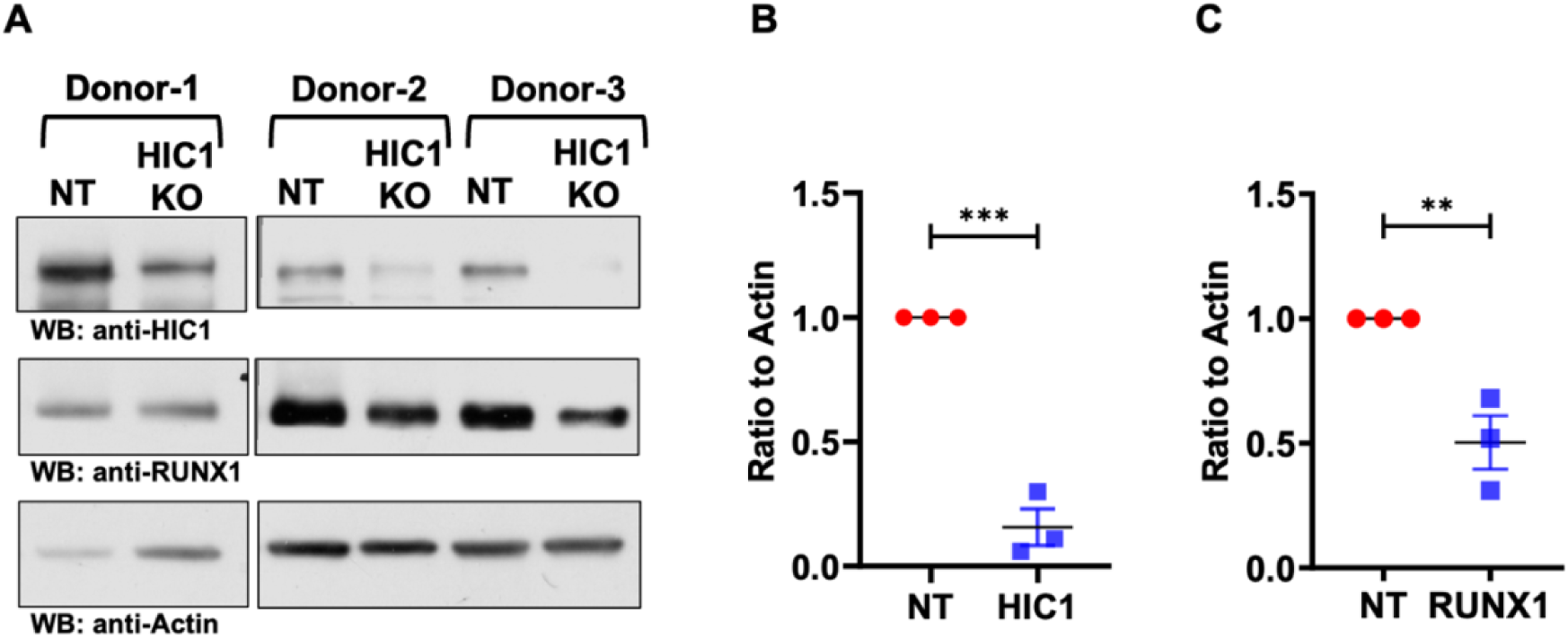
HIC1-knockout (KO) leads to reduced RUNX1 expression. (**A**) HIC1 was targeted with CRISPR-Cas9 RNPs in three individual donor cell pools, and HIC1, RUNX1 and Actin expression levels were measured by WB. (**B**) The dot plot shows the quantification of the HIC1 levels in HIC1 ablated and Control (NT) iTreg cells. (**C**) The dot plot shows the quantification of the RUNX1 levels in HIC1 silenced and Control (NT) iTreg cells. Quantification was performed using Image Studio Lite software Ver 5.2. HIC1 and RUNX1 levels were normalised to Actin levels. Statistical significance calculated using Student’s t-test (two-tailed paired) where ***denotes p value <0.0005. **denotes p value <0.005.

Collectively, these data suggest that HIC1 regulates RUNX1 expression in iTreg cells. RUNX1 is part of FOXP3 transcriptional complex and indispensable for Treg cell function [29,38]. Based on these findings, it is tempting to speculate that the reduction of RUNX1 expression in HIC1-deficient iTreg cells will lead to less RUNX1 availability, which may result in destabilization of the FOXP3-RUNX1-CBFB transcriptional complex and modulation of the FOXP3-RUNX1-CBFB complex dependent transcription program with increase in effector genes expression such as GATA3, TBX21 and IFNG with concomitant loss of suppression ability (Figure 5A, B).

**Figure 5:**
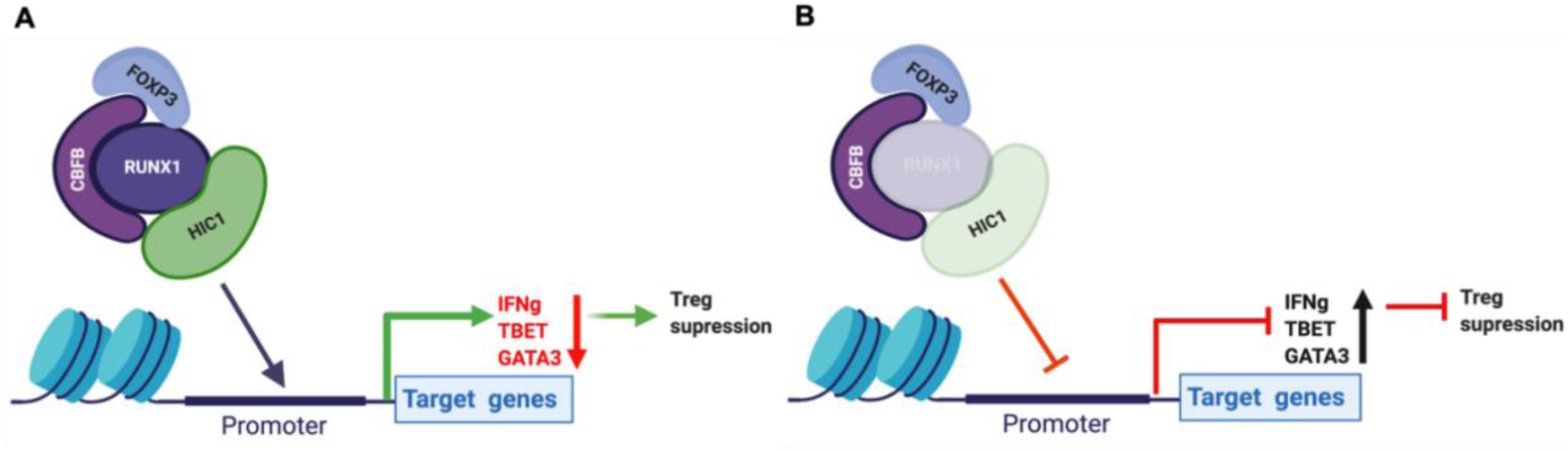
HIC1 interacts with FOXP3-RUNX1-CBFB protein complex. (**A**) FOXP3-RUNX1-CBFB-HIC1 protein complex regulates expression of TBET, IFNG and GATA3 and control Treg suppression. (**B**) HIC1 deficiency causes down-regulation of RUNX1, destabilizes the FOXP3 protein complex, increases the expression of effector T cell associated genes including TBET, GATA3 and IFNG which leads to loss of Treg suppression.

## 4. Discussion

Lineage specification factors play an essential role in Treg cell differentiation by regulating the expression of a set of genes that define the functional and phenotypic properties of Treg cells. Although FOXP3 is an important regulator of Treg cells, the differentiated status of Treg cells is not determined solely by FOXP3 expression and other factors play important role in this process. Recently, we established the importance of HIC1 in the development and suppressive function of iTreg cell [14]. The aim of this follow-up study was to elucidate the interactome of HIC1 in iTreg cells, and through this determined protein network gain mechanistic insights into how HIC1 regulates the development and suppressive capability of iTreg cells.

Notably, among the 61 high confidence HIC1 protein interactions identified, there were nine that are also known FOXP3 partners (i.e. BCLAF1, CBFB, DARS, DHX30, IKZF3, POLR2A, RBM8A and RUNX1) [37]. FOXP3 mediates the expression of its target genes through its association with a diverse set of binding partners [28,39]. This large complex includes transcriptional regulators and many sequence-specific TFs, e.g., RUNX1, NFAT, EOS, pSTAT3, IRF4, T-bet, GATA-3, RORγt, RORα, FOXO1 and FOXO3, SATB1 and HIF-1α [37,40].

RUNX1-CBFB heterodimer protein complex is part of the FOXP3 interacting protein network and play a pivotal role in the development and function of Treg cells [27]. Further, IKZF3 plays important role in epigenetic regulation by recruitment of chromatin modifiers such as the nucleosome remodeling and deacetylase (NuRD) complex to the locus of target genes to alter their chromatin organisation and hence their gene expression. IKZF3 interacts with FOXP3 to silence the transcriptional program of effector T cells and promote the differentiation of functional FOXP3^+^ Treg cells [28].

The results of this combination of bioanalytical methods, demonstrate that HIC1 is a part of FOXP3-RUNX1-CBFB transcription complex. This FOXP3 protein complex is indispensable for the expression of Treg signature genes and repressing the genes associated with effector functions, thus maintaining suppressive ability of iTreg cells [26–28,35]. Additionally, IKZF3 which is known to interact with FOXP3 and RUNX1 [37,41] was among the top proteins identified in the HIC1 interactome.

Our previous results indicated that upon HIC1 silencing, FOXP3 expression remained unchanged in iTreg cells [14]. Here, we demonstrate that HIC1 regulates the RUNX1 expression by binding to its promoter region. In the conditions of HIC1 deficiency, RUNX1 expression is reduced in iTreg cells. A lack of HIC1 and RUNX1 in HIC1 deficient cells may result in the destabilization of the FOXP3 transcriptional complex thus altering the FOXP3-RUNX1-CBFB dependent transcriptional program, which may in turn diminish Treg-specific gene expression and activate effector genes such as GATA3, TBX21 and IFNG with concomitant loss of suppression ability as observed in our earlier study [14].

Interestingly, we found BCOR as one of the top HIC1 interactors. BCOR is a transcriptional corepressor known to interact selectively with the BTB/POZ domain of the BCL6 transcriptional repressor which is involved in maintaining the lineage stability and regulating suppressive function of Treg cells [36,42]. The interaction between BCOR and BCL6 results in an increased transcriptional repression capacity of BCL6 [34]. Also, BCOR-mediated repression is important for Th17 cell differentiation [43]. Additional studies are needed to elucidate the role of BCOR - HIC1 interaction in context of iTreg cell differentiation and function.

ARS-interacting multi-functional proteins (AIMP1, AIMP2) were also detected as interactors of HIC1. Interestingly, it has been reported that AIMP1, enhance the differentiation of Treg cells, while it has no effect on Th1, Th2, and Th17 cell differentiation [44].

Further, among the identified HIC1 interactors were RPRD1B (CREPT), SRSF1, and HERC2 and PARP-1 (Figure 1A; Table S3). CREPT acts as an activator to promote transcriptional activity of the β-catenin-TCF4 complex in response to Wnt signaling in tumors [45]. Notably, it was reported that HIC1 also associates with β-catenin-TCF4 complex, and sequesters them to HIC1 bodies and modulates the Wnt signalling [46]. Wnt–β-catenin signaling is known to modulate function of human and murine Treg cells [47,48]. Based on these observations, HIC1 might contribute to Treg cell function by modulating Wnt–β-catenin signalling. However, further studies are needed to understand the role of association of HIC1 and CREPT in the context of Wnt–β-catenin signalling and iTreg cell differentiation.

SRSF1 is a member of the highly conserved serine/arginine (SR) family of RNA-binding proteins [49]. SRSF1 controls post-transcriptional gene expression via pre-mRNA alternative splicing, mRNA stability, and translation [32]. Recent studies show SRSF1 to be indispensable for Treg cell homeostasis and function in mouse [50].

HERC2 is a member of the HECT family of E3 ubiquitin-protein ligases and is implicated in DNA damage repair responses by ubiquitinating processes [33]. Ubiquitin-mediated processes influence the biology of Treg cells by modulating the signalling pathways critical for Foxp3 induction (e.g. TGFβ and NFκB signaling) or direct ubiquitination of the FOXP3 [51].

Poly (ADP-ribose) polymerase (PARP)-1 mediates PolyADP-ribosylation which plays a key role in the regulation of gene transcription in immune cells, including Treg cell differentiation [52,53]. It would be of immense interest to establish the role of HIC1interaction with SRSR1, HERC2 and PARP13 and how this association contributes to iTreg cell differentiation and function.

## 5. Conclusions

We present the first characterisation of the HIC1 interactome, which includes several noteworthy interactors. Gene ontology and pathway analysis of the associated proteins suggest that HIC1 is involved in several functions that have not been previously implicated. Our results indicate that HIC1 is a part of FOXP3 transcriptional complex comprised of RUNX1-CBFB and IKZF3. We propose that the compromised Treg suppression ability observed in HIC1 deficient iTreg cells may be partly due to the perturbed activity of the FOXP3 transcriptional complex.

## Author contributions

The conception and design of the study, or acquisition of data, or analysis and interpretation of data: SBAA, KB, OR, RM, SDB, IA, AM, TB, MHK, MMK, UUK, RL.

Drafting the article or revising it critically for important intellectual content: SBAA, KB, OR, RM, UUK, RL.

Final approval of the version to be submitted: SBAA, KB, OR, RM, SDB, TB, MHK, AM, IA, MMK, UUK, RL.

### Abbreviations

CBFB: Core-binding factor subunit beta
ChIPseq: Chromatin immunoprecipitation sequencing
FOXP3: Forkhead box P3
GO: Gene ontology
HIC1: Hypermethylated in cancer 1
IP: Immunoprecipitation
MS: Mass spectrometry
PLA: Proximity ligation assay
RUNX1: Runt-related transcription factor 1
SRM: Selected reaction monitoring
TF: Transcription factor
Treg: Regulatory T cells
iTreg: *In-vitro* induced Treg cells

## Supporting information

Supplementary tabls

## Acknowledgments

RL was supported by the Academy of Finland (AoF) Centre of Excellence in Molecular Systems Immunology and Physiology Research (2012-2017) grant 250114; by the AoF grants 292335, 294337, 292482, 319280, 329277, 331793, 335435 and 31444; by grants from the JDRF; the Novo Nordisk Foundation (grant NNF19OC0057218); the Sigrid Jusélius Foundation; Jane and Aatos Erkko Foundation and the Finnish Cancer Foundation. Our research is also supported by University of Turku, Åbo Akademi University, Turku Graduate School, InFLAMES Flagship Programme of the Academy of Finland (decision number: 337530). SBAA, was supported by InFLAMES Postdoctoral fellowship Programme, Finnish Cultural Foundation and Maud Kuistila Memorial Foundation. KB was supported by Orion Research Foundation sr and Finnish Cultural Foundation.

---

**Figure S1:**
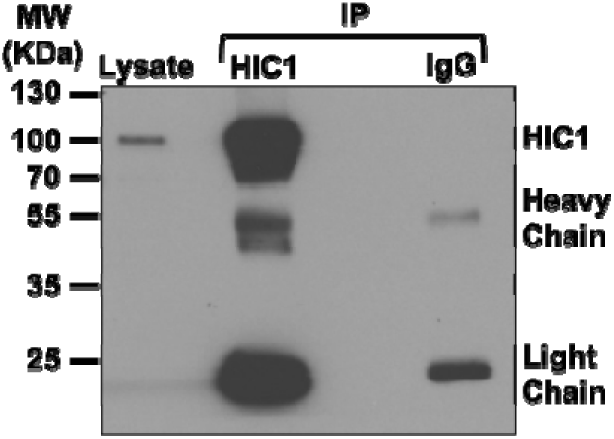
Naïve CD4^+^ CD25^-^ T cells were isolated from human umbilical cord blood and polarized to Treg lineage for 72 h. The cultured, Treg cells were lysed and HIC1 protein was immunoprecipitated (IP) using HIC1 specific antibody. Western blot (WB) analysis of HIC-1 immunoprecipitation (IP) with antibody specific to C-terminus region of HIC1, in 72 h polarized Treg cells. For HIC1 targeting antibody, an IgG control antibody was used. The figure is a representative WB of three experiments. Input, IgG control-IP and HIC1-IP reactions are shown.

